# Genome Report: A highly contiguous reference genome for Northern Bobwhite (*Colinus virginianus*)

**DOI:** 10.1101/729863

**Authors:** Jessie F. Salter, Oscar Johnson, Norman J. Stafford, William F. Herrin, Darren Schilling, Cody Cedotal, Robb T. Brumfield, Brant C. Faircloth

## Abstract

Northern bobwhites (*Colinus virginianus*) are small quails in the New World Quail family (Odontophoridae) and are one of the most phenotypically diverse avian species. Despite extensive research on bobwhite ecology, genomic studies investigating the evolution of phenotypic diversity in this species are lacking. Here, we present a new, highly contiguous assembly for bobwhites using tissue samples from a vouchered, wild, female bird collected in Louisiana. Using Dovetail Chicago and HiC libraries with the HiRise assembly pipeline, we produced an 866.8 Mbp assembly including 1,512 scaffolds with a contig N50 of 66.8 Mbp, a scaffold L50 of four, and a BUSCO completeness score of 90.8%. This new assembly greatly improves scaffold lengths and contiguity compared to an existing draft bobwhite genome and provides an important tool for future studies of evolutionary and functional genomics in bobwhites.

## INTRODUCTION

Northern bobwhites (*Colinus virginianus;* hereafter bobwhites) are widely distributed quails primarily found in pine woodlands and grasslands of the eastern United States and Mexico (Brennan 1999). Bobwhites hold a significant place in the cultural heritage of both countries due to their status as popular game birds (Bent 1963; Burger *et al.* 1999), and they have also played a significant role in biological research because they are one of the most intensively studied birds in the world (Guthery 1997). Bobwhites are remarkably polytypic: there are 22 subspecies recognized by male plumage (Brennan 1999) – a larger number of subspecies than 99% of all other birds (Dickinson and Remsen 2013). Although bobwhite ecology research has been extensive, the evolutionary relationships between bobwhite subspecies remain murky (Ellsworth *et al.* 1989; Evans *et al.* 2009; Eo *et al.* 2009; Williford 2013; Williford *et al.* 2014, 2016) and the genetic basis of phenotypic diversity in bobwhites has been largely unstudied (but see Cole *et al.* 1949).

Identifying genotypes associated with specific phenotypes increasingly relies on whole genome sequencing, particularly for investigating the genetic basis of phenotypic differences in non-model organisms (Ellegren 2014). The first draft genome assembly for bobwhites (GCA_000599485.1; hereafter Cv_TX_1.1) was generated from an unvouchered wild female bird from Texas (Halley *et al.* 2014). Cv_TX_1.1 used small and medium insert paired-end (PE) and mate pair (MP) libraries to produce a 1.172 Gb genome assembly with 77x coverage, 50% of the assembly in scaffolds of at least 45.5 Kbp in size (N50), and 90% of the assembly in 25,837 scaffolds (L90, Halley *et al.* 2014). Sequencing of additional PE and MP libraries from the same bird were used to generate a second assembly (GCA_000599465.2; hereafter Cv_TX_2.0), which yielded a 1.5-fold increase in coverage (122x), a 45-fold improvement in N50 (2.042 Mb), and a nearly 3-fold decrease in L90 (8,990 scaffolds; Oldeschulte *et al.* 2017). Although Cv_TX_2.0 was a marked improvement over Cv_TX_1.1, the scaffolds remained relatively short, which can hinder identification of structural variants (Domyan *et al.* 2014). Recent studies in birds and other taxa have demonstrated the importance of structural variants in generating morphological diversity within closely-related taxa (Lamichhaney *et al.* 2016; Tuttle *et al.* 2016; Vijay *et al.* 2016), highlighting the need for highly contiguous genome assemblies in phenotype-genotype studies (Wellenreuther and Bernatchez 2018).

Here, we describe Cv_LA_1.0, a new assembly for bobwhites using DNA extracted from a vouchered, wild female bird collected in Louisiana. To generate this assembly, we scaffolded contigs from small insert libraries with reads from Chicago (Putnam *et al.* 2016) and HiC (Lieberman-Aiden *et al.* 2009) methodologies and the HiRise assembly pipeline (Dovetail Genomics, LLC). The resulting Cv_LA_1.0 assembly is highly contiguous and represents a 32-fold increase in N50 and 528-fold decrease in L90 relative to Cv_TX_2.0 (Oldeschulte *et al.* 2017).

## METHODS

### Specimen collection and DNA extraction

We collected blood, liver, and other tissues for direct storage in liquid nitrogen from a wild, female bird legally harvested at Sandy Hollow Wildlife Management Area (30.827 N, 90.397 W) in Tangipahoa Parish, Louisiana. After tissue collection, we prepared a specimen for the LSU Museum of Natural Science (LSUMNS) Collection of Birds, and we stored tissue samples from this specimen in the LSUMNS Collection of Genetic Resources (LSUMZ B-91918). We shipped blood and liver to Dovetail Genomics, LLC (Scotts Valley, CA) where Dovetail Staff performed DNA extraction, library preparation, sequencing, and assembly steps. Dovetail staff extracted high molecular weight (HMW) DNA from tissues using the Blood and Cell Culture Midi Kit (Qiagen, GmbH) following the manufacturer’s protocol. Mean fragment length of the extracted DNA was 85 kb.

### Short-insert library preparation, sequencing, and assembly

Dovetail staff randomly fragmented extracted DNA by sonication using a Bioruptor Pico (Diagenode, Inc.) and 7 cycles of: sonication for 15 seconds followed by 90 seconds of rest. Dovetail staff then prepared a sequencing library by inputting fragmented DNA to the TruSeq DNA PCR-Free Library Preparation Kit (Illumina, Inc.) following the manufacturer’s protocol. Resulting libraries were sequenced on an Illumina HiSeq X platform using paired-end (PE) 150 bp sequencing. Resulting data were trimmed for low-quality bases and adapter contamination using Trimmomatic (Bolger *et al.* 2014) and used to assemble scaffolds with Meraculous v2.2.5 (Chapman *et al.* 2011). Before assembly, Dovetail staff used Jellyfish (Marçais and Kingsford 2011) with in-house software similar to GenomeScope (Vurture *et al.* 2017) to profile the short insert reads at a variety of k-mer values (25, 55, 85, 109), estimate genome size, and fit negative binomial models to the data. The resulting profiles suggested a k-mer size of 55 was optimal for assembly, and Dovetail staff assembled contigs using Meraculous with a k-mer size of 55, a minimum k-mer frequency of 12, and the diploid nonredundant haplotigs mode.

### Chicago library preparation and sequencing

Following *de novo* assembly with Meraculous, Dovetail staff prepared a single, proprietary “Chicago” library following the methods described in Putnam *et al.* (2016). Briefly, they reconstituted ~500 ng of HMW genomic DNA into chromatin *in vitro* and fixed the reconstituted DNA with formaldehyde. Then, they digested fixed chromatin with DpnII, filled in 5’ overhangs with biotinylated nucleotides, and ligated free, blunt ends. After ligation, they reversed crosslinks and purified the DNA from protein. Dovetail staff treated purified DNA to remove biotin that was not internal to ligated fragments and sheared the resulting DNA to ~350 bp mean fragment size using a Bioruptor Pico. Dovetail staff then prepared sequencing libraries from these sheared DNA using NEBNext Ultra enzymes (New England Biolabs, Inc.) and Illumina-compatible adapters. They isolated biotin-containing fragments using streptavidin beads before PCR enrichment of each library. Dovetail staff then sequenced amplified libraries on an Illumina HiSeq X platform using PE 150 reads.

### Dovetail HiC library preparation and sequencing (multiple libraries)

Dovetail staff also prepared two Dovetail HiC libraries following the procedures outlined in Lieberman-Aiden *et al.* (2009). Briefly, for each library, Dovetail staff used formaldehyde to fix chromatin in place in the nucleus. They extracted and digested fixed chromatin with DpnII, filled in the 5’ overhangs with biotinylated nucleotides, and ligated free blunt ends. After ligation, Dovetail staff reversed crosslinks and purified the DNA from protein. They treated the purified DNA to remove biotin that was not internal to ligated fragments and sheared the DNA to ~350 bp mean fragment size using a Bioruptor Pico. Dovetail staff then prepared sequencing libraries using NEBNext Ultra enzymes and Illumina-compatible adapters. They isolated biotin-containing fragments using streptavidin beads before PCR enrichment of each library and sequenced the resulting libraries on an Illumina HiSeq X Platform using PE 150 reads.

### Assembly scaffolding with HiRise

To scaffold and improve the bobwhite assembly, Dovetail staff input the *de novo* assembly from Meraculous, along with shotgun reads, Chicago library reads, and Dovetail HiC library reads into HiRise, a software pipeline designed for this purpose (Putnam *et al.* 2016). Using HiRise, Dovetail staff conducted an iterative analysis. First, they aligned shotgun and Chicago library sequences to the draft contig assembly using a modified SNAP read mapper (http://snap.cs.berkeley.edu). Second, they analyzed the separations of Chicago read pairs mapped within draft scaffolds to produce a likelihood model for genomic distance between read pairs, and they used this model to: identify and break putative misjoins, score prospective joins, and make joins above a threshold. Finally, after aligning and scaffolding the draft assembly using the Chicago data, Dovetail staff aligned and scaffolded the Chicago assembly using Dovetail HiC library sequences following the same method. After scaffolding, Dovetail staff used the short-insert sequences to close remaining gaps between contigs where possible.

### Assembly polishing, contiguity statistics, and BUSCO analyses

After receiving the assembly from Dovetail, we aligned the short insert data back to the scaffolded assembly using bwa v0.7.17-r1188 (Li and Durbin 2009) and samtools v1.9 (Li *et al.* 2009) and polished the scaffolds using Pilon v1.23 (Walker *et al.* 2014) on a 48-core, 1.5 TB RAM compute node with default parameters. After polishing, we computed contiguity statistics of our scaffolded assembly as well as the Cv_TX_2.0 assembly (Oldeschulte *et al.* 2017) using QUAST v5.0.2 (Mikheenko *et al.* 2018), UCSC Browser Utilities (Kent *et al.* 2002), and GNU Coreutils (https://www.gnu.org/software/coreutils), and we performed BUSCO analyses against both genomes using BUSCO v3.1.0 (Waterhouse *et al.* 2018) and the Aves Data Set (aves_odb9).

### Data availability

Data from all sequencing runs and the final assembly, Cv_LA_1.0, are available from NCBI BioProject (PRJNA454855). Short-insert, Chicago, and HiC reads are also available from the NCBI SRA (SRP215501), and the assembly is available from NCBI Genome using the accession VONY00000000. The version described in this manuscript is VONY01000000. Outputs from QUAST and BUSCO analyses are available from FigShare (doi: FILES SUBMITTED TO G3 FIGSHARE).

## RESULTS AND DISCUSSION

Sequencing of short-insert libraries produced 441.8 million read pairs with an average insert size of 428 bp. Analysis of the k-mer histogram at the optimal value of 55 suggested the genome size was 1.0 Mbp, and the estimated Q20 read depth for this genome size was approximately 118X. Meraculous assembly using a k-mer value of 55 produced 23,275 contigs having a total length of 853.1 Mbp and an N50 of 113.6 Kbp. These contigs were joined by Meraculous into 14,482 scaffolds totaling 854.1 Mbp in length with an N50 of 176.8 Kbp and a L90 of 5,343 scaffolds. The longest Meraculous scaffold was 1.6 Mbp. Meraculous estimated that the assembled contigs comprised 96% of the estimated, non-repetitive genome size and 84% of the entire genome size.

Chicago library sequencing produced 303 million read pairs, and the estimated physical coverage (the number of read pairs with inserts between 1 and 100 Kbp) spanning each position in the Meraculous assembly was 382.2. HiRise made 12,824 joins and one break to the Meraculous assembly to produce a Chicago assembly including 1,659 scaffolds totaling 866.68 Mbp in length with an N50 of 15.5 Mbp and a L90 of 53 scaffolds. The longest Chicago scaffold was 86 Mbp.

HiC library sequencing produced 111 million read pairs for Library 1 and 95 million read pairs for Library 2, and the estimated physical coverage (the number of read pairs with inserts between 10 and 10,000 kb) spanning each position in the Chicago assembly was 38,615. HiRise made 147 joins and zero breaks to the Chicago-scaffolded assembly to produce a HiC assembly including 1,512 scaffolds totaling 866.8 Mbp in length with an N50 of 66.9 Mbp and a L90 of 17 scaffolds. The longest HiC scaffold was 180.8 Mbp.

After polishing the HiC assembly, the bobwhite genome assembly Cv_LA_1.0 included 1,512 scaffolds having an N50 of 66.8 Mbp and a L50 of four (N90 = 13.1 Mbp; L90=17). Comparison of Cv_LA_1.0 with the Cv_TX_2.0 assembly (Table 1) shows the increase in contiguity of our assembly relative to the assembly produced by Oldeschulte *et al.* (2017). BUSCO analyses of both genomes are similar (Table 2), although we detected slightly fewer BUSCOs (-0.7%) in our Cv_LA_1.0 assembly relative to Cv_TX_2.0, perhaps due to repeat regions that were excluded from the contigs assembled by Meraculous.

**Table 1.** Metrics estimated using QUAST, UCSC Browser Utilities, and GNU Coreutils for *Colinus virginianus* genome assembly Cv_LA_1.0 (this manuscript) and comparison to a different assembly of a different individual, Cv_TX_2.0 (GCA_000599465.2; Oldeschulte *et al.* 2017), from the same species.

|  | **Cv_LA_1.0** | **Cv_TX_2.0** |
| --- | --- | --- |
| Contigs | 1,512 | 42,369 |
| Largest contig | 180,865,729 | 14,292,544 |
| Total length | 866,266,924 | 1,254,146,751 |
| N50 | 66,809,948 | 2,042,136 |
| N75 | 22,391,474 | 65,386 |
| N90 | 13,127,921 | 11,797 |
| L50 | 4 | 150 |
| L75 | 10 | 1,080 |
| L90 | 17 | 8,989 |
| GC (%) | 41.2 | 42.7 |
| # N's | 11,810,287 | 119,897,618 |
| # N's per 100 kbp | 1,363.4 | 9,560.1 |

**Table 2.** Genome completeness estimated using single copy orthologs (BUSCO v3) from *Colinus virginianus* assembly Cv_LA_1.0 (this manuscript) compared to a different assembly, Cv_TX_2.0 (GCA_000599465.2; Oldeschulte *et al.* 2017) from the same species.

|  | **Cv_LA_1.0** |  | **Cv_TX_2.0** |  |
| --- | --- | --- | --- | --- |
|  | count | percentage | count | percentage |
| Complete BUSCOs | 4,461 | 90.8% | 4,493 | 91.4% |
| Complete and single-copy BUSCOs | 4,416 | 89.8% | 4,435 | 90.2% |
| Complete and duplicated BUSCOs | 45 | 0.9% | 58 | 1.2% |
| Fragmented BUSCOs | 170 | 3.5% | 248 | 5.0% |
| Missing BUSCOs | 284 | 5.8% | 174 | 3.5% |
| Total BUSCO groups searched | 4,915 | - | 4,915 | - |

## ACKNOWLEDGEMENTS

We thank Shaune Hall and other staff members at Dovetail Genomics for working with us, and Donna Dittmann and Steve Cardiff for assistance with specimen preparation and tissue storage. Special thanks to Chick and Lulu for assistance in the field. Funding for this project was provided by Louisiana State University, National Science Foundation grants DEB-1242260 and IOS-1754417 (to BCF) and DEB-1655624 (to BCF and RTB). JFS was supported by the LSU Museum of Natural Science, and portions of this research were conducted with high performance computational resources provided by the Louisiana Optical Network Infrastructure (http://www.loni.org). The individual bobwhite used in this study was collected by a private quail hunter with a valid Louisiana hunting license. JFS and OJ prepared specimens, BCF performed analyses, JS and BCF wrote the paper, and BCF and RTB provided funding.

